# Changes of urinary proteome in rats after intragastric administration of calcium gluconate

**DOI:** 10.1101/2024.03.04.583150

**Authors:** Ziyun Shen, Minhui Yang, Haitong Wang, Youhe Gao

## Abstract

Calcium is an essential element for maintaining the normal physiological function of organisms. In this study, 3225 mg/kg/d calcium gluconate (equivalent to 300 mg/kg/d calcium) was intragastrically administered to rats for 4 days, and the urine proteome of rats was analyzed. Many differential proteins have been reported to be calcium related, such as Regucalcin (2.6 times higher after gavage than before gavage, p = 0.022), transmembrane protein 132A (8.2 times higher after gavage than before gavage, p = 0.009), creatine kinase (17.5 times higher before gavage than after gavage, p = 0.006), and claudin-3 (13.3 times higher before gavage than after gavage, p = 0.037). Differential protein enriched KEGG pathways included calcium signaling pathways, and biological processes and molecular functions also showed correlation with calcium. In this study, from the perspective of urine proteomics to explore the overall impact of calcium on the body, it is helpful to deeply understand the biological function of calcium and broaden the application potential of urine proteomics.

## 1 Introduction

Calcium is involved in a large number of important biological functions, is a key element in the maintenance of bone structure and cellular signaling, affects almost every aspect of cellular life, and is essential for the growth and development of the body^[1,2]^. Calcium is an essential element that can only be supplied through dietary sources. Calcium is an essential element that can only be supplied to the body through dietary sources. Current recommendations for dietary calcium range from 1000 to 1500 mg/d, depending on age and gender, and vary from country to country^[3]^. The amount recommended varies from country to country depending on age and gender.

Since urine is not part of the internal environment, in contrast to plasma, there is no mechanism for homeostasis, and it is able to accumulate early changes in the physiological state of the organism, reflecting more sensitively the changes in the organism, and is a source of next-generation biomarkers^[4]^. The proteins in urine contain a wealth of information that can reflect the small changes produced in different systems and organs of the organism.

Our laboratory has previously reported that the urine proteome is able to reflect the effects of magnesium threonate intake on the body in a more systematic and comprehensive manner, with the potential to provide clues for clinical nutrition research and practice^[5]^. The function of calcium as a key element in the maintenance of bone structure and cell signaling has been extensively studied. However, to date, there have been no studies exploring the overall effect of calcium on the organism from the perspective of the urinary proteome.

Calcium gluconate was chosen as a supplement in this study. Calcium gluconate is a common calcium supplement with high bioavailability and this form of calcium is widely used in the prevention and treatment of calcium deficiency. The aim of this study was to investigate the changes in the urinary proteome of rats after calcium gluconate intake, in the hope of deepening the understanding of the physiological functions of calcium, providing new perspectives for nutritional studies, and offering new clues for dietary regulation of human health and micronutrients.

## 2 Materials and Methods

### 2.1 Experimental materials

#### 2.1.1 Experimental consumables

5 ml sterile syringe (BD), gavage needle (16-gauge, 80 mm, curved needle), 1.5 ml/2 ml centrifuge tube (Axygen, USA), 50 ml/15 ml centrifuge tube (Corning, USA), 96-well cell culture plate (Corning, USA), 10 kD filter (Pall, USA), Oasis HLB solid phase extraction column (Waters, USA), 1ml/200ul/20ul pipette tips (Axygen, USA), BCA kit (Thermo Fisher Scientific, USA), high pH reverse peptide separation kit (Thermo Fisher Scientific, USA), iRT (indexed retention time, BioGnosis, UK).

#### 2.1.2 Experimental apparatus

Rat metabolic cages (Beijing Jiayuan Xingye Science and Technology Co., Ltd.), frozen high-speed centrifuge (Thermo Fisher Scientific, USA), vacuum concentrator (Thermo Fisher Scientific, USA), DK-S22 electric thermostatic water bath (Shanghai Jinghong Experimental Equipment Co., Ltd.), full-wavelength multifunctional enzyme labeling instrument (BMG Labtech, Germany), oscillator (Thermo Fisher Scientific, USA), TS100 constant temperature mixer (Hangzhou Ruicheng Instruments Co. BMG Labtech), electronic balance (METTLER TOLEDO, Switzerland), -80 □ ultralow-temperature freezing refrigerator (Thermo Fisher Scientific, USA), EASY-nLC1200 Ultra High Performance Liquid Chromatography system (Thermo Fisher Scientific, USA), and Orbitrap Fusion Lumos Tribird Mass Spectrometer (Thermo Fisher Scientific, USA) were used.

#### 2.1.3 Experimental reagents

Gluconate Calcium (Gluconate Calcium) was purchased from Shanghai Yuanye Biotechnology Co., Ltd, CAS No. 299-28-5, molecular formula C12H22CaO14, with a purity of more than 99%. In addition, Trypsin Golden (Promega, USA), dithiothreitol (DTT) (Sigma, Germany), iodoacetamide (IAA) (Sigma, Germany), ammonium bicarbonate NH4HCO3 (Sigma, Germany), urea (Sigma, Germany), and purified water (Wahaha, China) were used, Methanol for mass spectrometry (Thermo Fisher Scientific, USA), Acetonitrile for mass spectrometry (Thermo Fisher Scientific, USA), Purified water for mass spectrometry (Thermo Fisher Scientific, USA), Tris-Base (Promega, USA), Thiourea (Sigma, Germany) and other reagents.

#### 2.1.4 Analysis software

Proteome Discoverer (Version2.1, Thermo Fisher Scientific, USA), Spectronaut Pulsar (Biognosys, UK), Ingenuity Pathway Analysis (Qiagen, Germany); R studio (Version1.2.5001); Xftp 7; Xshell 7.

### 2.2 Experimental Methods

#### 2.2.1 Animal modeling

In this study, 16-week-old rats were used to minimize the effects of growth and development during gavage. Five healthy SD (Sprague Dawley) 9-week-old male rats (250±20 g) were purchased from Beijing Viton Lihua Laboratory Animal Technology Co. The rats were kept in a standard environment (room temperature (22±2)°C, humidity 65%-70%) for 7 weeks and weighed 500-600 g. The experiments were started, and all experimental operations followed the review and approval of the Ethics Committee of the School of Life Sciences, Beijing Normal University.

Dietary nutrient tolerable upper intake levels (UL): the average daily maximum intake of a nutrient for a population of a certain physiological stage and gender, which has no side effects or risks to the health of almost all individuals. Recommended nutrient intakes (RNI): intake levels that meet 97-98% of the individual needs of a particular age, gender, or physiological group.

According to the Dietary Guidelines for Chinese Residents, the tolerable upper intake levels (UL) of calcium is 2000mg/d. The UL for humans is converted to a dose of approximately 180mg/kg/d of calcium in rats based on body surface area and body weight^[6]^. In the present study, the dose of calcium in rats was 300 mg/kg/d by gavage, and the dose of calcium gluconate was 3225 mg/kg/d, which is 1.5 times of the UL in humans. 16.125g of calcium gluconate was dissolved in 500ml of sterile water and configured as a gavage solution. Each rat was gavaged with 5 ml of calcium gluconate solution once a day for 4 days. The first day of gavage was recorded as Ca-D1, and so on. Sampling time points were set before and after gavage for their own before and after control. The sample collected on the day before gavage was the control group, recorded as Ca-D0, and the sample numbered 21-25, and the sample collected on the 4th day of gavage was the experimental group, recorded as Ca-D4, and the sample numbered 31-35.

**Fig. 1.**
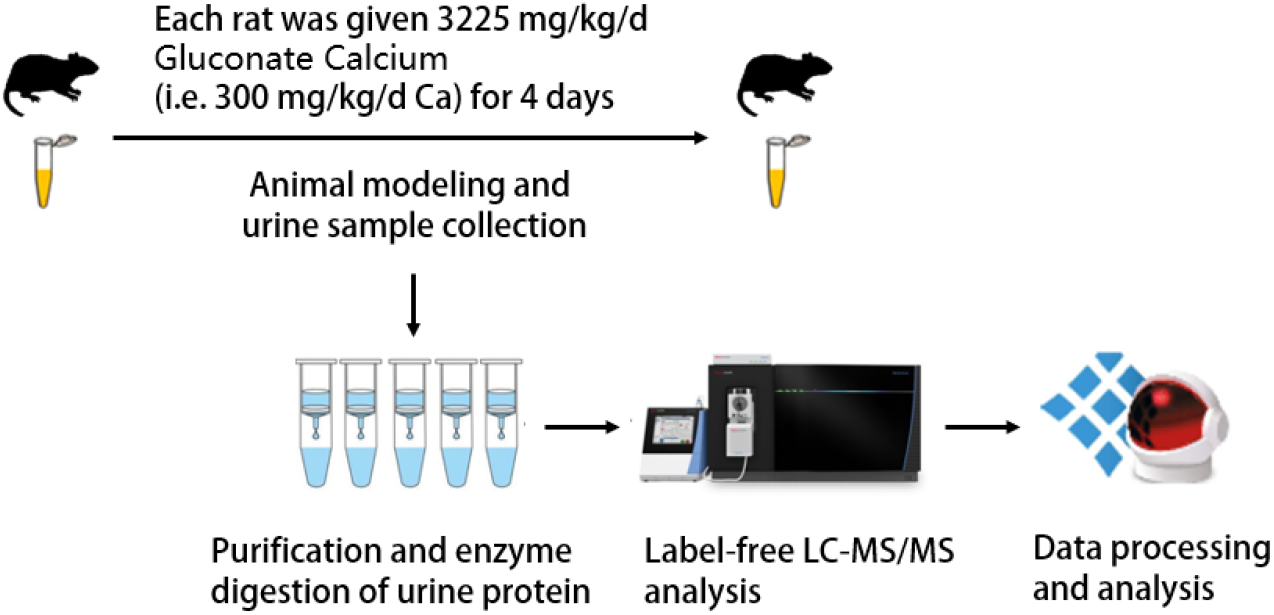
Research methodology and technical route

#### 2.2.2 Urine sample collection

One day before the start of gavage mineral supplementation (D0) and 4 days after gavage mineral supplementation (D4), each rat was individually placed in a metabolic cage at the same time, fasted and dehydrated for 12 h. Urine was collected overnight, and the urine samples were collected and placed in the refrigerator at -80°C for temporary storage.

#### 2.2.3 Urine sample processing

Two milliliters of urine sample was removed, thawed, and centrifuged at 4 °C and 12,000×g for 30 minutes. The supernatant was removed, and 1 M dithiothreitol (DTT, Sigma) storage solution (40 μl) was added to reach the working concentration of DTT (20 mM). The solution was mixed well and then heated in a metal bath at 37 °C for 60 minutes and allowed to cool to room temperature.

Then, iodoacetamide (IAA, Sigma) storage solution (100 μl) was added to reach the working concentration of IAM, mixed well and then reacted for 45 minutes at room temperature protected from light. At the end of the reaction, the samples were transferred to new centrifuge tubes, mixed thoroughly with three times the volume of precooled anhydrous ethanol, and placed in a freezer at -20 °C for 24 h to precipitate the proteins.

At the end of precipitation, the sample was centrifuged at 4 °C for 30 minutes at 10,000×g, the supernatant was discarded, the protein precipitate was dried, and 200 μl of 20 mM Tris solution was added to the protein precipitate to reconstitute it. After centrifugation, the supernatant was retained, and the protein concentration was determined by the Bradford method.

Using the filter-assisted sample preparation (FASP) method, urinary protein extracts were added to the filter membrane of a 10-kD ultrafiltration tube (Pall, Port Washington, NY, USA) and washed three times with 20 mM Tris solution. The protein was resolubilized by the addition of 30 mM Tris solution, and the protein was added in a proportional manner (urinary protein:trypsin = 50:1) to each sample. Trypsin (Trypsin Gold, Mass Spec Grade, Promega, Fitchburg, WI, USA) was used to digest proteins at 37 °C for 16 h.

The digested filtrate was the peptide mixture. The collected peptide mixture was desalted by an Oasis HLB solid phase extraction column, dried under vacuum, and stored at -80 °C. The peptide mixture was then extracted with a 0.1% peptide mixture. The lyophilized peptide powder was redissolved by adding 30 μL of 0.1% formic acid water, and then the peptide concentration was determined by using the BCA kit. The peptide concentration was diluted to 0.5 μg/μL, and 4 μL of each sample was removed as the mixed sample.

#### 2.2.4 LC-MS/MS analysis

All identification samples were added to a 100-fold dilution of iRT standard solution at a ratio of 20:1 sample:iRT, and the retention times were standardized. Data-independent acquisition (DIA) was performed on all samples, and each sample measurement was repeated 3 times, with 1-mix samples inserted after every 10 runs as a quality control. The 1-μg samples were separated using EASY-nLC1200 liquid chromatography (elution time: 90 min, gradient: mobile phase A: 0.1% formic acid, mobile phase B: 80% acetonitrile), the eluted peptides were entered into the Orbitrap Fusion Lumos Tribird mass spectrometer for analysis, and the corresponding raw files of the samples were generated.

#### 2.2.5 Data processing and analysis

The raw files collected in DIA mode were imported into Spectronaut software for analysis, and the highly reliable protein standard was peptide q value<0.01. The peak area quantification method was applied to quantify the protein by applying the peak area of all fragmented ion peaks of secondary peptides, and the automatic normalization was processed.

Proteins containing two or more specific peptides were retained, and missing values were replaced with 0. The amount of different proteins identified in each sample was calculated, and samples from rats before gavage of mineral supplements were compared with samples from rats 4 days after gavage of mineral supplements to screen for differential proteins.

Unsupervised cluster analysis (HCA), principal component analysis (PCA), and OPLS-DA analysis were performed using the Wukong platform (https://omicsolution.org/wkomics/main/). Functional enrichment analysis of differential proteins was performed using the DAVID database (https://david.ncifcrf.gov/) to obtain results in 3 areas: biological process, KEGG pathway and molecular function. Differential proteins and related pathways were searched based on Pubmed database (https://pubmed.ncbi.nlm.nih.gov/). Protein interaction network analysis was performed using the STRING database (https://cn.string-db.org/). Functions of differential proteins, Gene Ontology (GO) analysis results were searched using Uniprot database (https://www.uniprot.org/).

## 3 Results and discussion

### 3.1 Differential protein analysis

The missing values were replaced with 0. The pre-gavage samples of rats were compared with the samples on the 4th day of gavage and 63 differential proteins were screened. The conditions for screening differential proteins were: p-value <0.05 for t-test analysis, Fold change (FC) >2 or <0.5. as shown in Table 1.

**Table 1.** Differential proteins in the comparative analysis of the Ca-D0 and Ca-D4 groups (p-value <0.05, FC>2 or <0.5)

| From | Gene Names | FC | P | Related to Calcium |
| --- | --- | --- | --- | --- |
| Q3KRE3 | Gng10 | 0.041 | 0.0144 | [9,8,7] |
| D4A050 | Tbc1d32 | 0.041 | 0.040349 | [10] |
| A0A0G2K9J2 | Atp6v1h | 0.042 | 0.037623 | [11] |
| Q5BJT9 | Ckmt1 Ckmt1b | 0.057 | 0.00634 | [12] |
| Q63400 | Cldn3 | 0.075 | 0.036954 | [13] |
| A0A0G2JSG6 | Ak2 | 0.083 | 0.027132 |  |
| Q45QJ4 | Plcb3 | 0.086 | 0.049651 |  |
| P19139 | Csnk2a1 | 0.091 | 0.016627 | [14] |
| Q812E4 | Syt15 | 0.099 | 0.046653 | [15] |
| G3V9N7 | Pacsin3 | 0.126 | 0.036693 |  |
| A0A0G2JSV2 | cbr1;loc102556347;cbr1 | 0.134 | 0.036766 | [16] |
| B2GV72 | Cbr3 | 0.148 | 0.049133 | [16] |
| P13233 | Cnp | 0.169 | 0.040811 | [17] |
| A0A0G2JZY6 | Sptbn1 | 0.196 | 0.040082 | [18] |
| P28576 | Pdgfa Rpa1 | 0.201 | 0.028378 | [19] |
| A0A0G2JVT6 |  | 0.313 | 0.041395 |  |
| P30120 | Timp1 | 0.316 | 0.032539 |  |
| P31977 | Ezr | 0.442 | 0.045434 | [20] |
| F1MAD3 | Pkd1 | 2.001 | 0.001833 | [21] |
| Q3KRD8 | Eif6 Itgb4bp | 2.013 | 0.032475 | [22] |
| Q4V8K5 | Brox | 2.036 | 0.006032 |  |
| Q62867 | Ggh | 2.043 | 0.038097 | [23] |
| F1LNJ2 | Snrnp200 | 2.062 | 0.009847 |  |
| Q5I0D5 | Lhpp | 2.074 | 0.01541 | [24] |
| D3ZY96 | Ngp | 2.126 | 0.03711 |  |
| P53369 | Nudt1 Mth1 | 2.153 | 0.012753 | [25] |
| A0A0G2JZ40 | Reck | 2.163 | 0.015578 |  |
| Q68FS4 | Lap3 | 2.306 | 0.041286 |  |
| D3ZKN1 | Bpifa6 | 2.342 | 0.044608 |  |
| P85971 | Pgls | 2.366 | 0.012947 |  |
| Q9Z2L0 | Vdac1 | 2.397 | 0.047346 | [26] |
| P48508 | Gclm Glclr | 2.397 | 0.030914 |  |
| F7F1Y3 | Vps4a | 2.491 | 0.001948 |  |
| D3ZFC6 | Itih4 | 2.509 | 0.022924 | [27] |
| A0A0G2K7Y0 | Cr11 Cd46 | 2.551 | 0.024081 | [28] |
| D3ZGN2 | Cpne5 | 2.605 | 0.005304 | [29] |
| F1LLW8 | Ids | 2.609 | 0.001086 |  |
| D3ZZT9 | Col14a1 | 2.626 | 0.025971 | [30] |
| Q03336 | Rgn Smp30 | 2.647 | 0.02213 | [31] |
| P23928 | Cryab | 2.769 | 0.022525 | [32] |
| P19468 | Gclc | 2.833 | 0.041759 |  |
| D4A5U3 | Tgm3 Tgase3 | 3.029 | 0.002246 | [33] |
| P07379 | Pck1 | 3.042 | 0.014027 | [34] |
| Q9WUW9 | Sult1c2a Sultk2 | 3.149 | 0.04358 |  |
| G3V712 | Krt7 | 3.188 | 0.023808 | [35] |
| P23606 | Tgm1 | 3.516 | 0.037304 | [36] |
| Q5M872 | Dpep2 | 3.566 | 0.036296 |  |
| A0A0G2JSX3 | Naglt1 | 3.583 | 0.02385 | [37] |
| D3ZFH5 | Itih2 | 3.588 | 0.048854 | [38] |
| Q9ESG3 | Cltrn Nx17 Tmem27 | 3.847 | 0.021334 |  |
| Q6IFZ5 | Krt76 Kb9 | 3.848 | 0.006512 | [35] |
| A0A0G2JU92 | Emb | 4.051 | 7.58E-05 | [39] |
| D4A1A6 | Il36rn | 4.089 | 0.000233 | [40] |
| F1M7X4 | ErbB4 | 4.1 | 0.04026 | [41] |
| D3ZCT9 | Gal3st1 | 4.526 | 0.035962 |  |
| F1LX20 | Tnfaip8l3 | 4.601 | 0.005274 | [42] |
| D3ZDX5 | Plxnb1 | 4.936 | 0.04226 | [43] |
| Q7TQ70 | Fga | 5.237 | 0.038916 | [44] |
| A0A0G2JUF6 | Idh2 | 5.434 | 0.044302 | [45] |
| Q5BJY9 | Krt18 | 5.807 | 0.041656 | [35] |
| P81155 | Vdac2 | 6.22 | 0.03652 | [46] |
| D3ZVB6 | Lpal2 LOC690326 | 6.556 | 0.014939 | [47] |
| Q80WF4 | Tmem132a Gbp Hspa5bp1 | 8.158 | 0.008658 | [48] |

The 63 differential proteins were analyzed for protein function and literature search using PubMed database to analyze the relationship between protein and calcium in detail one by one. This was done by using the pubmed database and entering the protein name of the differential proteins into the search box along with calcium, with the search range being title/abstract, e.g., “Calcium [Title/Abstract] AND Protein [Title/Abstract]”. The literature was then read to confirm the relationship between the differential proteins and calcium. Also, the functions of the differential proteins were analyzed according to the Uniprot database.

Regucalcin (RC) is a calcium-binding protein that is capable of regulating calcium ion signaling, calcium ion-dependent cellular processes, and enzyme activities^[31]^, and has been implicated in biological processes such as intracellular calcium ion homeostasis and regulation of calcium-mediated signaling.

Polycystin 1 (PC1), voltage-dependent anion-selective channel protein 1 (VDAC1) has functions such as calcium ion transmembrane transport, calcium channel activity, etc. PC1 is a regulator of calcium ion permeable cation channel signaling receptor^[21]^. PC1 is a signaling receptor that regulates calcium-permeable cation channels. VDAC1 expression is up-regulated by elevated calcium ion levels^[26]^. VDAC1 and VDAC2 are differential proteins, and the voltage-dependent anion channel (VDAC) acts as a channel for the exchange of ions (including Ca2+) across the outer mitochondrial membrane^[49]^. The FC for VDAC2 was 6.2 with a P value of 0.037.

The functions of protein kinase C and casein kinase substrate in neurons 3 include calcium channel inhibitor activity, negative regulation of calcium ion transport.

The molecular functions of 1-phosphatidylinositol 4,5-bisphosphate phosphodiesteras include binding to calcium ion and calmodulin binding.

Iduronate 2-sulfatase, protein-glutamine gamma-glutamyltransferase E (also known as glutamyltransferase E, Transglutaminase E,TGE) function includes binding to calcium ions. Glutamine transferase is a widely distributed calcium-dependent protease that is widely distributed^[33]^.

Copine 5 is a calcium-dependent lipid-binding intracellular protein^[29]^. The functions of Copine 5 include calcium-dependent phospholipid binding, cellular response to calcium ions.

Collectrin (a.k.a. Transmembrane protein 27) is involved in biological processes including calcium-regulated cytokinesis.

Creatine kinase (FC=0.057, p=0.006) regulates sodium-calcium exchanger activity^[12]^ .

The FC for the guanine nucleotide-binding protein subunit is 0.041 and the p-value is 0.014. signaling through guanine nucleotide-binding proteins is associated with cellular calcium homeostasis^[9,8,7]^ .

Claudin-3 (FC=0.075, p=0.037) mediates paracellular transport of cations such as calcium ions^[13]^.

Receptor protein-tyrosine kinase (FC=4.1,p=0.04) is able to interact with calmodulin^[41]^ that are involved in the calcium signaling pathway.

TNF alpha induced protein 8 like 3 had an FC of 4.6 with a p-value of 0.005. Interleukin-1 (IL-1) had an FC of 4.1 with a p-value of 0.0002. Elevated intracellular calcium ions lead to the production of pro-inflammatory cytokines, including IL-1β and tumor necrosis factor-α (TNF-α)^[42]^.

Transmembrane protein 132A (TMEM132A) had an FC of 8.2 and a p-value of 0.009. treatment with calcium modulators significantly attenuated TMEM132A mRNA and protein levels in NSC-132 cells^[48]^.

Due to space limitations, other literature showing differential protein-calcium correlations are listed in Table 1.

### 3.2 Biological pathway analysis

The 157 differential proteins (P value <0.05, FC>2 or <0.5) were imported into the DAVID database and enriched to 22 biological processes (BPs) (as shown in Table 2). 8 differential proteins were enriched to aging as a biological process, and 9 differential proteins were enriched to negative regulation of apoptotic processes. Four differentially differentiated proteins were enriched for the biological processes of keratinization, response to insulin, positive regulation of protein kinase B signaling, kidney development, actin cytoskeleton organization, and positive regulation of ERK1 and ERK2 cascades, respectively. Three differential proteins were enriched for liver regeneration, positive regulation of phosphatidylinositol 3-kinase signaling, and other biological processes, respectively. Most of the differential proteins enriched to biological pathways were reported to be associated with calcium. The literature showing the relevance of biological processes to calcium is listed in Table 2.

**Table 2.** Biological processes (BP) to which Ca-D0 and Ca-D4 groups were enriched for differential proteins (p-value < 0.05)

| Term | Count | % | P-Value | Related to Calcium |
| --- | --- | --- | --- | --- |
| aging | 8 | 12.9 | 8.10E-05 | [50] |
| negative regulation of apoptotic process | 9 | 14.5 | 5.90E-04 | [51] |
| keratinization | 4 | 6.5 | 1.30E-03 | [52] |
| regulation of protein localization to plasma membrane | 3 | 4.8 | 5.00E-03 |  |
| digestive tract development | 3 | 4.8 | 8.80E-03 |  |
| response to insulin | 4 | 6.5 | 9.00E-03 | [53] |
| cellular response to thyroxine stimulus | 2 | 3.2 | 9.00E-03 | [54] |
| cysteine metabolic process | 2 | 3.2 | 9.00E-03 |  |
| response to nitrosative stress | 2 | 3.2 | 1.20E-02 | [55] |
| regulation of mitochondrial depolarization | 2 | 3.2 | 1.50E-02 | [56] |
| positive regulation of memory T cell differentiation | 2 | 3.2 | 2.10E-02 | [57] |
| positive regulation of phosphatidylinositol 3-kinase signaling | 3 | 4.8 | 2.10E-02 | [58] |
| positive regulation of protein kinase B signaling | 4 | 6.5 | 2.20E-02 | [58] |
| kidney development | 4 | 6.5 | 2.30E-02 | [59] |
| liver regeneration | 3 | 4.8 | 2.50E-02 | [60] |
| hyaluronan metabolic process | 2 | 3.2 | 3.30E-02 |  |
| glutathione biosynthetic process | 2 | 3.2 | 3.30E-02 | [61] |
| actin cytoskeleton organization | 4 | 6.5 | 3.40E-02 | [62] |
| response to human chorionic gonadotropin | 2 | 3.2 | 3.80E-02 | [64,63] |
| positive regulation of ERK1 and ERK2 cascade | 4 | 6.5 | 3.90E-02 | [65] |
| glutamate metabolic process | 2 | 3.2 | 4.10E-02 | [66] |
| response to activity | 3 | 4.8 | 4.90E-02 |  |

### 3.3 Molecular function and KEGG pathway analysis

The 157 differential proteins (P value <0.05, FC >1.5 or <0.67) were imported into the DAVID database and enriched to 10 molecular functions (Table 3), including structural molecule activity, glutamate-cysteine ligase activity, kinase activity, voltage-gated anion channel activity, macromolecular complex binding, ceramide binding, protein-glutamine γ-glutamyltransferase activity, carbonyl reductase (NADPH) activity, metallopeptide endonuclease inhibitor activity, and porin activity.

**Table 3.** Molecular functions (MF) enriched to differential proteins in Ca-D0 and Ca-D4 groups (p-value < 0.05)

| Term | Count | % | P-Value |
| --- | --- | --- | --- |
| structural molecule activity | 6 | 9.7 | 8.90E-05 |
| glutamate-cysteine ligase activity | 2 | 3.2 | 6.40E-03 |
| kinase activity | 4 | 6.5 | 1.30E-02 |
| voltage-gated anion channel activity | 2 | 3.2 | 1.60E-02 |
| macromolecular complex binding | 7 | 11.3 | 2.20E-02 |
| ceramide binding | 2 | 3.2 | 2.80E-02 |
| protein-glutamine gamma-glutamyltransferase activity | 2 | 3.2 | 2.80E-02 |
| carbonyl reductase (NADPH) activity | 2 | 3.2 | 4.10E-02 |
| metalloendopeptidase inhibitor activity | 2 | 3.2 | 4.70E-02 |
| porin activity | 2 | 3.2 | 4.70E-02 |

The 157 differential proteins (P value <0.05, FC >1.5 or <0.67) were imported into the DAVID database and enriched to six KEGG pathways (Table 4), including glutathione metabolism, cofactor biosynthesis, metabolic pathways, neutrophil extracellular trap formation, iron death, and calcium signaling pathways.

**Table 4.** KEGG pathways enriched to differential proteins in Ca-D0 and Ca-D4 groups (P value <0.05)

| Term | Count | % | P-Value |
| --- | --- | --- | --- |
| Glutathione metabolism | 4 | 6.5 | 2.40E-03 |
| Biosynthesis of cofactors | 5 | 8.1 | 2.50E-03 |
| Metabolic pathways | 14 | 22.6 | 4.10E-03 |
| Neutrophil extracellular trap formation | 5 | 8.1 | 5.00E-03 |
| Ferroptosis | 3 | 4.8 | 1.10E-02 |
| Calcium signaling pathway | 5 | 8.1 | 1.40E-02 |

Among them, five differential proteins were enriched in the calcium signaling pathway, including platelet-derived growth factor subunit A (PDGF-1), voltage-dependent anion-selective channel protein 1 (VDAC-1), 1-phosphatidylinositol 4,5-bisphosphate, 1-phosphatidylinositol 4,5-bisphosphate phosphodiesterase, voltage-dependent anion-selective channel protein 2 (VDAC-2), and receptor protein-tyrosine kinase.

## 4 Perspective

The results illustrate that short-term supplementation of calcium gluconate affects the organism, and the urine proteome of rats can show changes in calcium-related proteins and biological functions, and also suggests that the urine proteome can reflect the overall changes of the organism comprehensively and systematically. The present study provides clues for the in-depth understanding of the metabolic process, mechanism of action, and biological function of calcium in organisms from the perspective of urine proteomics, as well as provides new research perspectives for future related studies.

